# Integrative genomics of the mammalian alveolar macrophage response to intracellular mycobacteria

**DOI:** 10.1101/2020.08.25.266668

**Authors:** Thomas J. Hall, Michael P. Mullen, Gillian P. McHugo, Kate E. Killick, Siobhán C. Ring, Donagh P. Berry, Carolina N. Correia, John A. Browne, Stephen V. Gordon, David E. MacHugh

**Author notes:** Genuity Science, Cherrywood Business Park. Loughlinstown, Co. Dublin, D18 K7W4, Ireland. Correspondence and requests for materials should be addressed to D.E.M.

## Abstract

Bovine tuberculosis (bTB), caused by infection with *Mycobacterium bovis*, is a major disease affecting cattle globally as well as being a zoonotic risk to human health. The key innate immune cell that first encounters *M. bovis* is the alveolar macrophage, previously shown to be substantially reprogrammed during intracellular infection by the pathogen. Here we use multi-omics and network biology approaches to analyse the macrophage transcriptional response to *M. bovis* infection and identify core infection response pathways and gene modules. These outputs were integrated with results from genome-wide associations of *M. bovis* infection to enhance the detection of putative genomic variants for disease resistance. Our results show that network-based integration of relevant transcriptomics data can extract additional information from large genome-wide associations and that this approach could also be used to integrate relevant functional genomics outputs with results from genomic association studies for human tuberculosis caused by the related *Mycobacterium tuberculosis*.

## Introduction

Bovine tuberculosis (bTB) is a chronic disease of livestock, particularly among domestic dairy and beef cattle, which has been conservatively estimated to cause more than $3 billion annual losses to global agriculture (Steele 1995; Waters *et al.* 2012). The disease can also establish across a large variety of wildlife species including, for example, American bison (*Bison bison*), African buffalo (*Syncerus caffer*), the brushtail possum (*Trichosurus vulpecula*), red deer (*Cervus elaphus*), wild boar (*Sus scrofa*) and the European badger (*Meles meles;* (Fitzgerald & Kaneene 2013; Palmer 2013; Malone & Gordon 2017; Gormley & Corner 2018). The aetiological agent of bTB, *Mycobacterium bovis*, is a member of the *Mycobacterium tuberculosis* complex (MTBC) and has a genome sequence 99.95% identical to *M. tuberculosis*, the primary cause of human tuberculosis (TB) (Garnier *et al.* 2003). *M. tuberculosis* is a highly successful pathogen that is the leading cause of human deaths from a single infectious agent—approximately 1.25 million in 2018 (World Health Organization 2019). In addition, for several low-and middle-income countries, the human TB disease burden is increased by zoonotic TB (zTB) caused by infection with *M. bovis* (Thoen *et al.* 2016; Olea-Popelka *et al.* 2017; Vayr *et al.* 2018; Luciano & Roess 2020).

Scientific understanding of bTB and human TB has been synergistically intertwined since the 19^th^ century and the foundational research work of Theobald Smith and others (Daniel 2006; Cambau & Drancourt 2014; Malone & Gordon 2017). The pathogenesis of bTB disease in cattle is comparable with human TB disease and many aspects of *M. bovis* infection are also characteristic of *M. tuberculosis* infection (Neill *et al.* 2001; Russell 2003; Cassidy 2006; Pollock *et al.* 2006; Waters *et al.* 2014). Consequently, it is now widely recognised that *M. bovis* infection of cattle and bTB disease represent an important comparative system for understanding human TB caused by *M. tuberculosis* (Hein & Griebel 2003; Van Rhijn *et al.* 2008; Waters *et al.* 2011; Williams & Orme 2016; Gong *et al.* 2020).

Inhalation of aerosolized bacteria is the main route of transmission for *M. bovis* in cattle and the primary site of infection is normally the lungs (Palmer *et al.* 2002; Cassidy 2006; Palmer *et al.* 2019). Here the bacilli are phagocytosed by alveolar macrophages (AM)—key effector cells of the innate immune system, which provide surveillance of pulmonary surfaces and can normally destroy or restrict inhaled intracellular bacilli (Weiss & Schaible 2015; Kaufmann & Dorhoi 2016). *M. bovis* and other facultative intracellular MTBC pathogens have evolved a complex range of mechanisms to evade, subvert, and exploit innate immune responses, thereby facilitating colonisation, persistence and replication within host macrophages (de Chastellier 2009; Cambier *et al.* 2014; Schorey & Schlesinger 2016; Awuh & Flo 2017). These mechanisms include: recruitment of cell surface receptors on the host macrophage through molecular mimicry; restricting phagosome maturation and autophagy; detoxification of reactive oxygen species and reactive nitrogen intermediates (ROSs and RNIs); modulation of type I interferon (IFN) signalling; suppression of antigen presentation; rewiring and short-circuiting of macrophage signal transduction pathways; manipulation of host macrophage metabolism; egress of bacilli into the macrophage cytosol; and inhibition of apoptosis with concomitant induction of necrosis leading to immunopathology and shedding by the host to complete the pathogenic life cycle (Hussain Bhat & Mukhopadhyay 2015; Queval *et al.* 2017; BoseDasgupta & Pieters 2018; Chaurasiya 2018; Stutz *et al.* 2018; Leopold Wager *et al.* 2019; Russell *et al.* 2019). Hence after infection, a two-way response is triggered between the pathogen and macrophage, the outcome of which ultimately leads to establishment of infection or clearance of the pathogen. The latter outcome of clearance may, or may not, require engagement of the adaptive immune system. As the detection of *M. bovis* infection in cattle generally relies on detecting an adaptive immune response to the pathogen, the outcome of which is slaughter of positive animals (‘reactors’), identifying genes that underpin efficacious innate responses promises to reveal favourable genomic variants for incorporation into breeding programmes.

Since 2005, substantial efforts have been made to better understand host-pathogen interaction for bTB using transcriptomics technologies such as gene expression microarrays and RNA sequencing (RNA-seq) at the host cellular level—specifically the bovine AM (bAM) and initial innate immune responses to infection by *M. bovis* (Widdison *et al.* 2008; Widdison *et al.* 2011; Magee *et al.* 2014; Nalpas *et al.* 2015; Vegh *et al.* 2015; Malone *et al.* 2018; Hall *et al.* 2020). These studies have helped to define a “pathogenic signature” (Falkow 2008; Cambier *et al.* 2014) of *M. bovis* infection in bAM, which reflects the tension between macrophage responses to contain and kill intracellular pathogens and evasion and avoidance mechanisms evolved by these mycobacteria. Using functional genomics data mining of transcriptomics data, it has also been shown that bAM responses to *M. bovis* infection can be clearly differentiated from infection with *M. tuberculosis*, the primary cause of human TB (Malone *et al.* 2018). In addition, these studies have been expanded to encompass surveys of the bAM epigenome using methylome sequencing and chromatin immunoprecipitation sequencing (ChIP-seq). This work has demonstrated that the transcriptional reprogramming of bAM caused by *M. bovis* infection is profoundly shaped by chromatin remodelling at gene loci associated with critical components of host-pathogen interaction (O’Doherty *et al.* 2019; Hall *et al.* 2020).

In parallel to functional genomics studies of bTB, increasingly powerful genome-wide association studies (GWAS) have been performed in Irish and UK cattle populations using estimated breeding values (EBVs; estimate of genetic merit of an animal derived from a statistical model) for several *M. bovis* infection resistance traits with heritabilities ranging from 0.04 to 0.37, depending on the phenotype used (Finlay *et al.* 2012; Bermingham *et al.* 2014; Richardson *et al.* 2016; Raphaka *et al.* 2017; Wilkinson *et al.* 2017; Tsairidou *et al.* 2018b; Ring *et al.* 2019). These GWAS have used medium-and high-density single-nucleotide polymorphism (SNP) arrays and, more recently, imputed whole-genome sequence (WGS) data sets for a large multi-breed GWAS on 7,346 bulls, which identified 64 quantitative trait loci (QTLs) associated with resistance to *M. bovis* infection (Ring *et al.* 2019); the association study was based on phenotypic data from 781,270 individuals.

We have recently shown that integration of bAM functional genomics data sets—RNA-seq, microRNA-seq and ChIP-seq—with a GWAS data set for resistance to *M. bovis* infection can be used to enhance detection of genomic regions associated with reduced incidence of bTB disease (Hall *et al.* 2020). For the present study, we substantially expand this work by leveraging gene-focused network-and pathway-based methods under a statistical framework based on a new software tool, *gwinteR*, to integrate transcriptomics data from *M. bovis*-challenged bAM with WGS-based GWAS results for resistance to *M. bovis* infection (Ring *et al.* 2019). The primary aim of this work is to evaluate whether this approach can systematically enhance detection of genomic sequence variants and genes underpinning bTB disease resistance in cattle populations.

## Methods

### Animal Ethics

All animal procedures were performed in accordance with EU Directive 2010/63/EU and Irish Statutory Instrument 543/2012, with ethical approval from the University College Dublin (UCD) Animal Research Ethics Committee (AREC-13–14-Gordon).

### Genomics Data Acquisition and Computational and Bioinformatics Workflow

Genome-wide RNA-seq transcriptomics data from a 48-h bAM time course challenge experiment using the sequenced *M. bovis* AF2122/97 strain have been previously generated by our group (GEO accession: GSE62506). The complete laboratory methods used to isolate, culture and infect bAM with *M. bovis* AF2122/9 and generate strand-specific RNA-seq libraries using RNA harvested from these cells are described in detail elsewhere (Magee *et al.* 2014; Nalpas *et al.* 2015; Malone *et al.* 2018). Briefly, these RNA-seq data were generated using bAM obtained by lung lavage of ten unrelated age-matched 7 to 12-week-old male Holstein-Friesian calves. Bovine AM were either infected *in vitro* with *M. bovis* AF2122/97 or incubated with media only. Following total RNA extraction from *M. bovis*-infected and control non-infected alveolar macrophages, 78 strand-specific RNA-seq libraries were prepared (paired-end 2 × 90 nucleotide reads). These comprised *M. bovis*-and non-infected samples from each post-infection time point (2, 6, 24 and 48 hours post-infection [hpi]) across 10 animals with the exception of one animal that did not yield sufficient alveolar macrophages for *in vitro* infection at 48 hpi.

GWAS data sets for the present study were obtained from intra-breed imputed WGS-based GWAS analyses that used estimated breeding values (EBVs) derived from a bTB infection phenotype, which were generated for 2,039 Charolais, 1,964 Limousin and 1,502 Holstein-Friesian sires (Ring *et al.* 2019). The bTB phenotype, the WGS-based imputed SNP data, and the quantitative genetics methods are described in detail elsewhere (Ring *et al.* 2019); however, the following provides a brief summary. The bTB infection phenotype was defined for every animal present during each herd-level bTB breakdown when a bTB reactor or a slaughterhouse case was identified. Cattle that yielded a positive single intradermal comparative tuberculin test (SICTT), and/or post-mortem lymph node lesion, or laboratory culture result/s were coded as bTB = 1 and all other cattle present in the herd during the bTB-breakdown were coded as bTB = 0; potential exposure of cattle within the bTB breakdown was also considered in this study (Ring *et al.* 2019). After phenotype data edits, bTB resistance EBVs were generated for 781,270 phenotyped cattle (plus their recorded ancestors). After within-breed SNP filtering using thresholds for minor allele frequency (MAF ≤ 0.002) and deviation from Hardy-Weinberg equilibrium (HWE; *P* < 1 × 10^-6^), there were 17,250,600, 17,267,260 and 15,017,692 autosomal SNPs for the 2,039 Charolais (CHA), 1,964 Limousin (LIM) and 1,502 Holstein-Friesian (HOFR) sire analyses. A single-SNP regression analyses was performed for each breed separately using weighted (i.e., by an effective record contribution) sire EBVs for *M. bovis* infection resistance/susceptibility and the nominal *P*-values were used for downstream integrative genomics analyses.

All data-intensive computational procedures were performed on a 36-core/72-thread compute server (2× Intel^®^ Xeon^®^ CPU E5-2697 v4 processors, 2.30 GHz with 18 cores each), with 512 GB of RAM, 96 TB SAS storage (12 × 8 TB at 7200 rpm), 480 GB SSD storage, and with Ubuntu Linux OS (version 18.04 LTS). The complete computational and bioinformatics workflow is available with additional information as a public GitHub repository (github.com/ThomasHall1688/Bovine_multi-omic_integration). The individual components of the experimental and computational workflows are shown in Fig. 1 and described in more detail below.

**Fig. 1.**
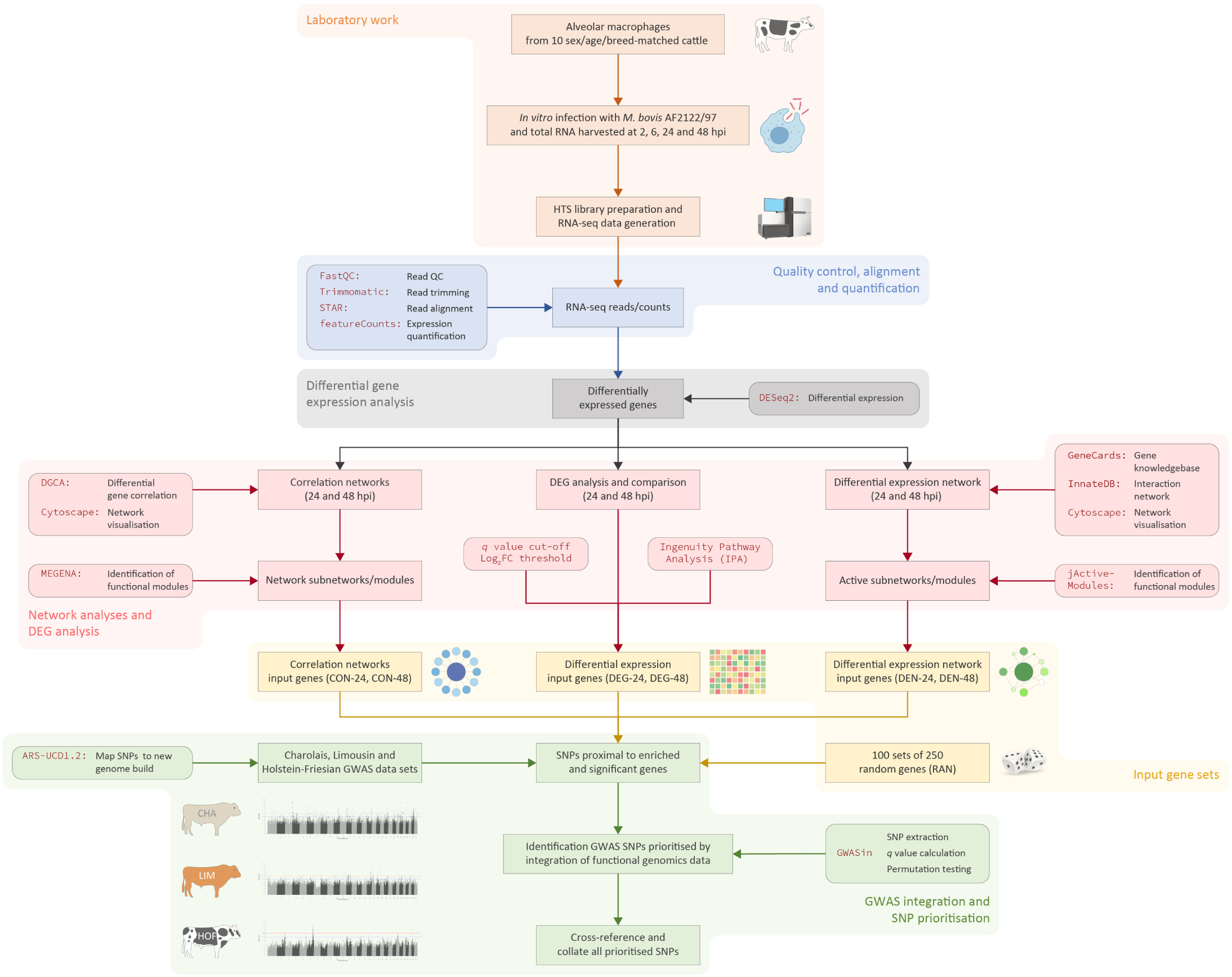
Schematic showing the experimental and computational workflow use to integrate bAM transcriptomics outputs and *M. bovis* infection resistance trait GWAS data

### Differential gene expression analysis of RNA-seq data

A custom Perl script was used to deconvolute barcoded RNA-seq reads into individual libraries, filter out adapter sequence reads, and remove poor quality reads (Nalpas *et al.* 2015). At each stage of the process, a quality check was performed on the FASTQ files with FastQC (version 0.11.8) (Andrews 2019). Paired-end sequence reads were then aligned to the *Bos taurus* reference genome (ARS-UCD1.2, GenBank assembly accession: GCA_002263795.2) (Rosen *et al.* 2020) using the STAR aligner (version 2.7) (Dobin *et al.* 2013). Read counts for each gene were calculated using featureCounts (version 1.6.4) (Liao *et al.* 2014), set to unambiguously assign uniquely aligned paired-end reads in a stranded manner to gene exon annotation. Using the R statistical programming language (version 3.5.3) (R Core Team 2019), gene annotation was derived from the NCBI database via a GFF annotation file GCF_002263805.1 with additional descriptions and chromosomal locations annotated using GO.db (version 3.8.2) (Carlson 2019) and biomaRt packages (version 2.40.0) (Durinck *et al.* 2009). Differential gene expression analysis was performed using the DESeq2 package (version 1.24.0) (Love *et al.* 2014) with a longitudinal time series design that accounted for time (hpi) and experimental treatment (*M. bovis*-infected versus control). Lowly expressed reads were removed using the mean of normalized counts as a filter statistic; individual genes with very low read counts would typically not exhibit significant differential expression due to high dispersion (Love *et al.* 2014). In addition, extreme count outliers were removed within DESeq2 using the Cook’s distance (Cook 1977) as previously described (Love *et al.* 2014). Multiple testing correction was performed on each time point using the Benjamini-Hochberg (B-H) false discovery rate (FDR) method (Benjamini & Hochberg 1995). The default criteria for differentially expressed (DE) genes were an FDR-adjusted *P*-value less than 0.05 (*P*_adj_. < 0.05) and an absolute log_2_ fold-change greater than one (|log_2_FC| > 1).

### Ingenuity^®^ Pathway Analysis (IPA) of differentially expressed genes

Ingenuity^®^ Pathway Analysis—IPA^®^ (version 1.1, summer 2020 release; Qiagen, Redwood City, CA, USA) was used to perform a statistical enrichment analysis of DE gene sets and expression data (Kramer *et al.* 2014). This enabled identification of canonical pathways and functional processes of biological importance in alveolar macrophages challenged with *M. bovis* across the longitudinal infection time course. Following best practice, the default background gene set for pathway and functional process enrichment testing was the set of detectable genes across all RNA-seq libraries for each time point contrast and not the complete bovine transcriptome (Timmons *et al.* 2015).

### Identification of functional gene modules using differential co-expression network analysis

An integrated computational pipeline for differential co-expression network analysis was implemented using: 1) the Differential Gene Correlation Analysis (DGCA) R package (version 1.0.2) (McKenzie *et al.* 2016); 2) the Cytoscape open source Java platform for visualisation and integration of biomolecular interaction networks (version 3.7.0) (Shannon *et al.* 2003); and 3) the Multiscale Embedded Gene Co-expression Network Analysis (MEGENA) R package (version 1.3.7) (Song & Zhang 2015).

Due to the low variance in gene expression observed between samples at 2 and 6 hpi (Fig. 1b), only data from the 24 and 48 hpi time points were used for the differential correlation analysis. The DGCA R package was used to filter normalised gene counts such that genes in the lower 30th percentile of median expression values were removed. Pearson correlation coefficients were then calculated for each gene pair between the control non-infected and *M. bovis*-infected samples at 24 and 48 hpi. Following this, for each time point, the infected and control samples were randomly shuffled, and the analysis was repeated for a total of ten iterations. Additionally, using the permutation testing and reference pool distribution approaches implemented in DGCA, an empirical *P*-value was calculated for each observed correlation coefficient and *q*-values were calculated based on empirical *P*-values and the estimated proportion of null hypotheses; gene pairs were then considered to be differentially correlated with a *q*-value threshold of 0.10 (McKenzie *et al.* 2016).

The correlation networks and network parameters generated by DGCA were initially visualised, examined and evaluated using the Cytoscape platform. In correlation networks, based on gene co-expression, each gene acts as a node and each correlation acts as a weighted edge, depending on the strength of the correlation coefficient (Schaefer *et al.* 2017; van Dam *et al.* 2018). The DGCA network data for the 24 and 48 hpi time points were then imported into MEGENA for identification of functional subnetworks (modules) using differentially correlated (*q* < 0.10) gene pairs ranked by *q*-value to a maximum of 3,500 gene pairs; this gave 3,500 and 1,085 gene pairs for the 24 and 48 hpi correlation networks, respectively. Each gene pair was assigned to a class that described the change in correlation depending on infection status, and only those genes that exhibited a change in expression pattern were included in the MEGENA analysis to identify functional modules.

Functional modules in the 24 and 48 hpi correlation networks were detected as locally coherent subnetwork clusters with a minimum of 20 unique genes that MEGENA classified as statistically significant (*P*-value < 0.05) based on analyses of shortest path indices, local path index, weight of the correlation and overall modularity. The resulting MEGENA functional modules were visualised using the Cytoscape platform and genes embedded in functional modules at 24 and 48 hpi were combined and annotated for downstream GWAS integration as described below. The genes contained in these functional modules were also subject to gene ontology (GO) term enrichment analyses within the MEGENA package (Song & Zhang 2015).

### Detection of active gene subnetworks using a tuberculosis and mycobacterial infection gene interaction network

The GeneCards^®^ gene compendium and knowledgebase (http://www.genecards.org; version 4.9), which integrates multiple sources of biological information on all annotated and predicted human genes (Stelzer *et al.* 2016), was used to identify a set of genes that are functionally associated with the host response to TB and other diseases caused by infection with mycobacteria. The search query used was `tuberculosis OR mycobacterium OR mycobacteria OR mycobacterial` and genes were ranked by a GeneCards statistic—the *Relevance Score*—based on the *Elasticsearch* algorithm (Gormley & Tong 2015), which determines the strength of the relationships between genes and keyword terms. Gene IDs were converted to human Ensembl gene IDs (Yates *et al.* 2020) and retained for downstream analysis using the InnateDB knowledgebase and analysis platform for systems level analysis of the innate immune response (http://www.innatedb.com; version 5.4) (Breuer *et al.* 2013).

A gene interaction network (GIN) was generated with the gene list output from GeneCards using InnateDB with default settings and this network was visualised using Cytoscape. The jActivesModules Cytoscape plugin (version 3.12.1) (Ideker *et al.* 2002) was then used to superimpose the bAM RNA-seq gene expression data and detect, through a greedy search algorithm, differentially active subnetworks (modules) of genes at the 24 and 48 hpi time points. Locally coherent clusters that contain genes that are differentially expressed were identified using the log_2_FC and *P*_adj_. values of each differentially expressed gene; the overall connectivity of those genes with their immediate module co-members; and the comparison of that connectivity with a background comprised of randomly drawn networks using the same genes, but independent of the base network. Genes embedded in active modules that were detected as statistically significant at 24 and 48 hpi were combined and annotated for downstream GWAS integration as described below.

### Integration of *M. bovis*-infected bovine AM gene expression data with bTB GWAS data

To facilitate integration of GWAS data with gene sets generated from functional genomics data analyses, an R software package was developed—*gwinteR* (github.com/ThomasHall1688/gwinteR), which can be used to test the hypothesis that a specific set of genes is enriched for signal in a GWAS data set relative to the genomic background. This gene set, for example, could be an output from an active gene module network analysis of transcriptomics data from a cell type or tissue relevant to the GWAS phenotype. To formally test the primary hypothesis, the *gwinteR* tool was designed to determine if genomic regions containing GWAS SNPs that are proximate to genes within a gene set are enriched for statistical associations with the trait/s analysed in the GWAS.

The *gwinteR* tool works as follows: 1) a set of significant and non-significant SNPs (named the target SNP set) is collated across all genes in a specific gene set at increasing genomic intervals upstream and downstream from each gene inclusive of the coding sequence (e.g., ±10 kb, ±20 kb, ±30 kb… …±100 kb); 2) for each genomic region, a null distribution of 1,000 SNP sets, each of which contains the same number of total significant and non-significant combined SNPs as the target SNP set, is generated by resampling with replacement from the search space of the total population of SNPs in the GWAS data set; 3) the nominal (uncorrected) GWAS *P*-values for the target SNP set and the null distribution SNP sets are converted to local FDR-adjusted *P*-values (*P*_adj_.) using the fdrtool R package (current version 1.2.15) (Strimmer 2008); 4) a permuted *P*-value (*P*_perm_.) to the test the primary hypothesis for each observed genomic interval target SNP set is generated based on the proportion of permuted random SNP sets where the same or a larger number of SNPs exhibiting significant *q*-values (e.g. *q* < 0.05 or *q* < 0.10) are observed; 5) *gwinteR* generates data to plot *P*_perm_. results by genomic interval class and obtain a graphical representation of the GWAS signal surrounding genes within the target gene set; 6) a summary output file of all SNPs in the observed target SNP set with genomic locations and *q*-values is generated for subsequent investigation.

In the original bTB GWAS data set used for the present study (Ring *et al.* 2019), the WGS-imputed SNPs were mapped to the UMD3.1 bovine genome assembly (Zimin *et al.* 2009). Consequently, prior to GWAS data integration, the imputed and previously filtered SNPs for each of the three breed groups were mapped, using a custom R pipeline (github.com/ThomasHall1688/Bovine_multi-omic_integration), to the most recent ARS-UCD1.2 cattle reference genome assembly (Rosen *et al.* 2020). After this step, there were 14,583,567, 14,586,972 and 12,740,315 autosomal SNPs with nominal GWAS *P*-values that could be used for integrative genomics analyses of the CHA, LIM and HOFR breeds, respectively.

For the integrative analyses of bAM functional genomics outputs with the bTB GWAS data, three different subsets of genes were used: 1) basic DE gene sets that were filtered to ensure manageable computational loads using stringent expression threshold criteria of |log_2_FC| > 2 and *P*_adj_. < 0.01 and *P*_adj_. < 0.000001 for 24 and 48 hpi, respectively; 2) genes embedded in functional modules at 24 and 48 hpi that were detected using the MEGENA package in the differential co-expression network analyses; and 3) genes embedded in active modules at 24 and 48 hpi that were identified using jActiveModules within the tuberculosis and mycobacterial infection interaction network.

## Results

### Differential gene expression and pathway analyses of *M. bovis*-infected bovine AM

Quality filtering of RNA-seq read pairs yielded a mean of 22,681,828 ± 3,508,710 reads per individual library (*n* = 78 libraries). A mean of 19,582,959 ± 3,021,333 read pairs (86.17%) were uniquely mapped to locations in the ARS-UCD1.2 bovine genome assembly. Detailed filtering and mapping statistics are shown in Supplementary Data 1 and multivariate PCA analysis of the individual animal sample expression data using DESeq2 revealed separation of the control and *M. bovis*-infected bAM groups at the 24 and 48 hpi time points, but not at the 2 and 6 hpi time points (Supplementary Figure 1).

Using default criteria for differential expression (FDR *P*_adj_. < 0.05; |log_2_FC| > 1), and considering the *M. bovis-*infected bAM relative to the control non-infected macrophages, three DE genes were detected at 2 hpi (all three exhibited increased expression in the *M. bovis*-infected group); 97 DE genes were detected at 6 hpi (40 increased and 57 decreased); 1,345 were detected at 24 hpi (764 increased and 581 decreased); and 2,915 were detected at 48 hpi (1,528 increased and 1,387 decreased) (Fig. 2a; Supplementary Data 2). Fig. 2b shows that 2,982 genes were differentially expressed across the 24 and 48 hpi time points. Table 1 shows a breakdown of DE genes across the infection time course for a range of statistical thresholds and fold-change cut-offs, including the default criteria (FDR *P*_adj_. < 0.05; |log_2_FC| > 1). To ensure manageable computational loads, the DE gene sets that were used for GWAS integration with *gwinteR* were filtered with |log_2_FC| > 2, and *P*_adj_. < 0.01 and *P*_adj_. < 0.000001 for 24 and 48 hpi, respectively. With these criteria, there were 378 input genes for GWAS integration identified at 24 hpi and 390 input genes at 48 hpi. (24 hpi and 48 hpi DEG gene sets). In addition, 210 genes overlapped between the two time points. The two DEG *gwinteR* input gene sets (DEG-24 and DEG-48 – see Fig. 1) are also detailed in Supplementary Data 2.

**Table 1:**
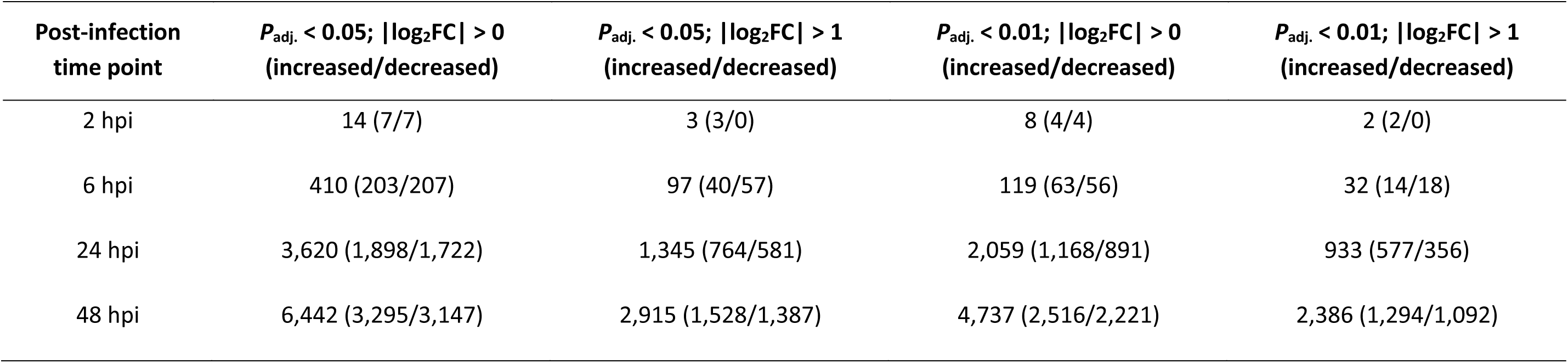
Differentially expressed genes detected in *M. bovis*-infected bovine AM relevant to control non-infected bAM

**Fig. 2.**
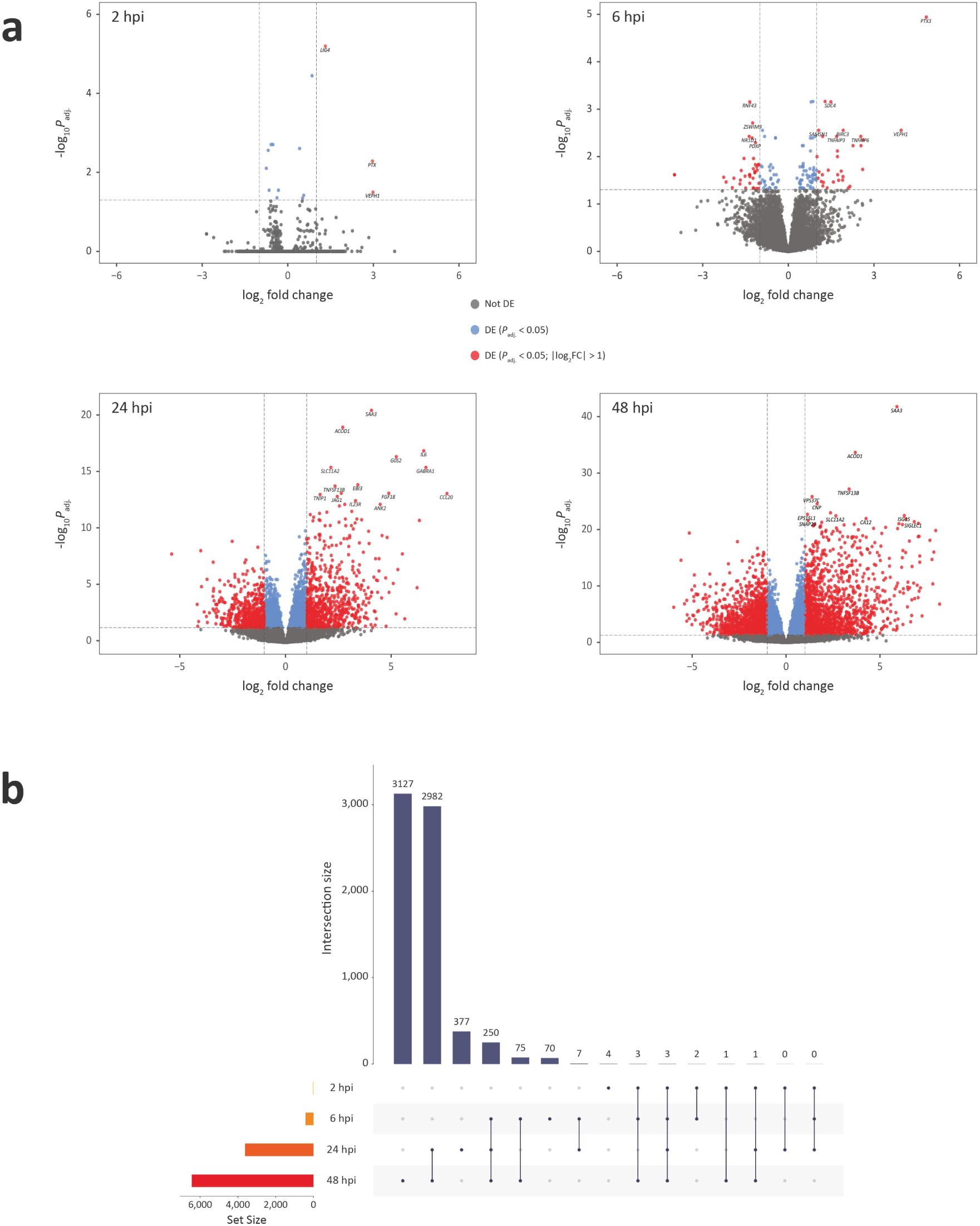
Differentially expressed genes in *M. bovis*-infected bAM at 2, 6, 24, and 48 hpi. **a** Volcano plots of differentially expressed genes with FDR *P*_adj_. value thresholds of 0.05 and absolute log_2_ fold-change > 1. **b** UpSet plot showing the intersection of shared differentially expressed genes across the four post-infection time points.

To produce gene sets for the IPA Core Analysis within the recommended range for the number of input entities (Kramer *et al.* 2014; Qiagen 2019) and to include DE genes with small fold-change values, gene sets were filtered using only *P*_adj_. thresholds of 0.05 and 0.01 at 24 hpi and 48 hpi, respectively. In addition, the target species selected was *Homo sapiens* and the cell type used was *Macrophage* with the *Experimentally Observed* and *High Predicted* confidence settings. This resulted in 1,957 input genes (1,071 upregulated and 886 downregulated) from a background detectable set of 16,084 at 24 hpi and 2,492 input genes (1,401 upregulated and 1,091 downregulated) from a background detectable set of 17,492 genes at 48 hpi. The IPA analysis was focused on the 24 and 48 hpi time points because a relatively small numbers of DE genes were detected at 2 and 6 hpi (Fig. 2 and Table 1).

Using the B-H method for multiple test correction in IPA (*P*_adj_. < 0.05), there were 68 and 48 statistically significant enriched IPA canonical pathways at 24 hpi and 48 hpi, respectively (Supplementary Data 3). Enriched pathways at 24 hpi included *Role of Pattern Recognition Receptors in Recognition of Bacteria and Viruses*, *IL-6 Signalling*, *TNFR2 Signalling*, *Role of RIG1-like Receptors in Antiviral Innate Immunity*, *Role of Cytokines in Mediating Communication between Immune Cells*, *Communication between Innate and Adaptive Immune Cells*, *IL-12 Signalling and Production in Macrophages*, *IL-10 Signalling*, *Protein Ubiquitination Pathway*, *Toll-like Receptor Signalling*, *NF-κB Signalling*, *PI3K/AKT Signalling*, and *TNFR1 Signalling*. The most highly activated pathway at 24 hpi was *PI3K/AKT Signalling*. Enriched pathways at 48 hpi included *Protein Ubiquitination Pathway*, *Role of Cytokines in Mediating Communication between Immune Cells*, *IL-12 Signalling and Production in Macrophages*, *Role of RIG1-like Receptors in Antiviral Innate Immunity*, *Role of Pattern Recognition Receptors in Recognition of Bacteria and Viruses*, *Communication between Innate and Adaptive Immune Cells*, *TNFR2 Signalling*, *Role of PI3K/AKT Signalling in the Pathogenesis of Influenza*, *IL-10 Signalling*, and *Toll-like Receptor Signalling*. The SIGORA software tool (Foroushani *et al.* 2013) has been previously used to identify biological pathways associated with a robust ‘core’ bAM response to infection with both *M. bovis* and *M*. *tuberculosis* (Malone *et al.* 2018). It is therefore reassuring that many of these pathways—including *PI3K-Akt Signalling Pathway*, *RIG-I-like Receptor Signalling Pathway*, *Toll-like Receptor Signalling* and *Protein Ubiquitination Pathway*—were also enriched using the IPA methodology at 24 and 48 hpi.

### Differential co-expression correlation networks and identification of functional gene modules

For the generation of bAM differential co-expression correlation networks, filtering of genes with low measure of central tendency, which reduces the number of potential spurious correlations (McKenzie *et al.* 2016), resulted in 11,354 and 11,170 genes at 24 and 48 hpi, respectively. Following this step, differential correlation analysis using DGCA with an empirical *P*_adj_. value threshold of 0.10 resulted in 3,507 differentially correlated gene pairs out of 128,913,316 total pairwise correlations at 24 hpi; and 1,135 from a total of 124,768,900 at 48 hpi (Supplementary Data 4). The correlation networks generated at 24 hpi and 48 hpi (Fig. 3a) yielded a total of 22 and 14 functional gene modules, respectively (Fig. 3b and Supplementary Data 4). After removal of duplicates, consolidated totals of 460 genes and 416 genes were contained in the functional modules at 24 hpi and 48 hpi, respectively. There were also 26 genes that overlapped between the functional modules for the two time points. The two correlation network (CON) *gwinteR* input gene sets (CON-24 and CON-48 – see Fig. 1) are also detailed in Supplementary Data 4.

**Fig. 3.**
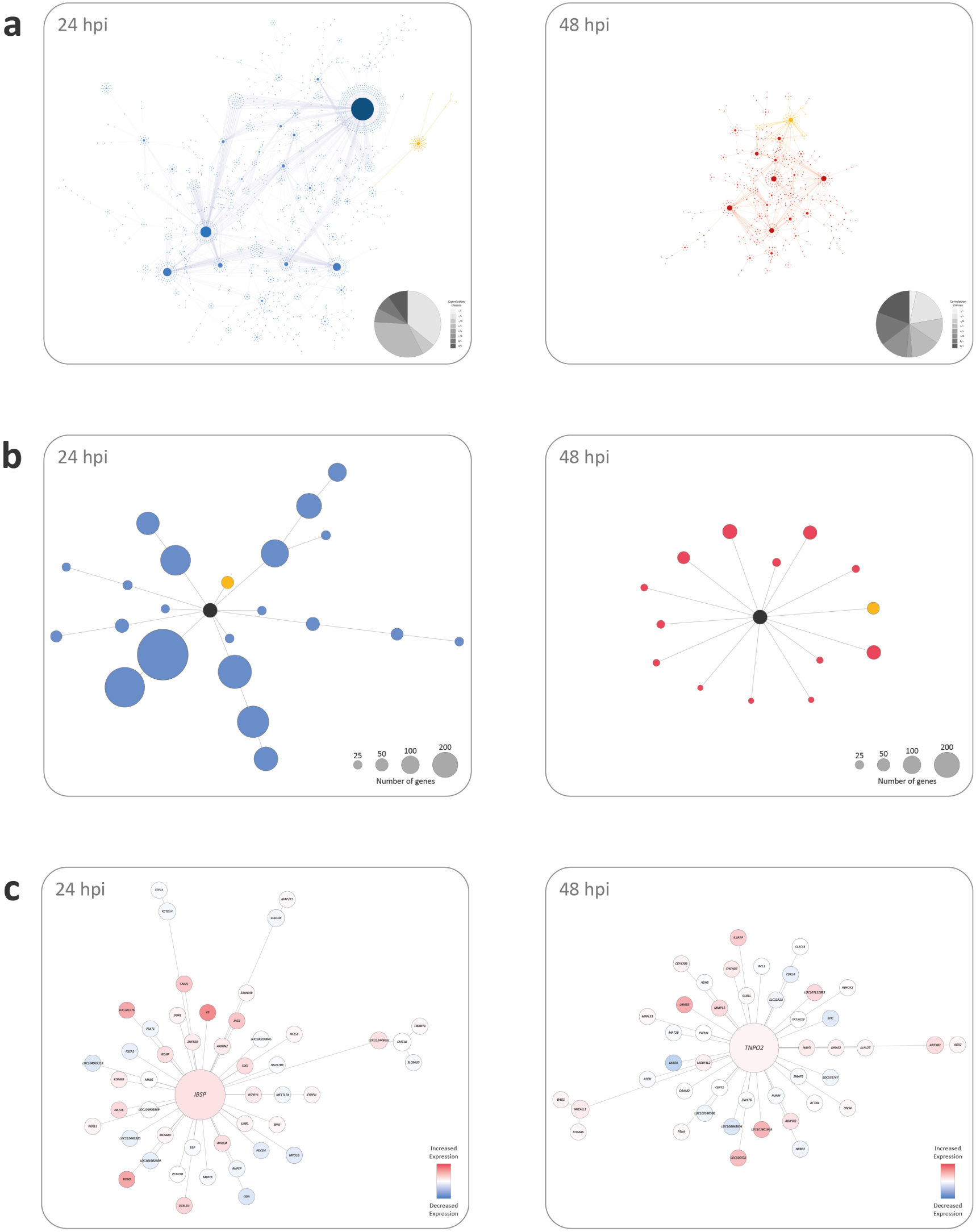
Differential co-expression correlation networks and submodules at 24 and 48 hpi. **a** The complete correlation networks for *M. bovis*-infected bAM at 24 and 48 hpi. **b** The subnetwork modules detected at 24 hpi and the 14 subnetwork modules detected at 48 hpi with individual example modules highlighted in yellow. **c** Individual example subnetwork modules at 24 and 48 hpi.

GO term enrichment was also performed for each functional module at 24 hpi and 48 hpi, with the top three GO terms retained for each functional module (Supplementary Figure 2 and Supplementary Figure 3). The top five GO terms at 24 hpi (ranked by *P*_adj_.) were *translation* (GO:0006412), *peptide biosynthetic process* (GO:0043043), *amide biosynthetic process* (GO:0043604), *structural constituent of ribosome* (GO:0003735), and *cellular amide metabolic process* (GO:0043603) (Supplementary Figure 2). The top five enriched GO terms at 48 hpi (ranked by *P*_adj_.) were *signalling receptor activity* (GO:0038023) and *molecular transducer activity* (GO:0060089), *transforming growth factor beta activation* (GO:0036363), *chemokine activity* (GO:0008009), and *signalling receptor binding* (GO:0005102) (Supplementary Figure 3).

### Differential expression network analysis and identification of activated modular subnetworks

The GeneCards search query generated a total of 2,291 gene hits (Supplementary Data 5) using the search terms: `tuberculosis OR mycobacterium OR mycobacteria OR mycobacterial`. To provide a computationally manageable number of genes for an InnateDB input data set (Breuer *et al.* 2013), a GeneCards *Relevance Score* (GCRS) threshold was used (GCRS > 2.5). This GCRS cut-off produced an input list of 258 functionally prioritised genes for generation of an InnateDB gene interaction network (GIN) and the top ten genes from this list ranked by GCRS were: interferon gamma receptor 1 (*IFNGR1*), interleukin 12 receptor subunit beta 1 (*IL12RB1*), toll like receptor 2 (*TLR2*), solute carrier family 11 member 1 (*SLC11A1*), signal transducer and activator of transcription 1 (*STAT1*), interleukin 12B (*IL12B*), cytochrome b-245 beta chain (*CYBB*), tumour necrosis factor (*TNF*), interferon gamma receptor 2 (*IFNGR2*), and interferon gamma (*IFNG*).

The large GIN produced by InnateDB starting with the input list of 258 functionally prioritised genes was visualised using Cytoscape and consisted of 7,001 nodes (individual genes) and 19,713 edges (gene interactions) (Fig. 4a). Supplementary Data 5 provides information for all gene interactions represented in Fig. 4a. Following visualisation of the large GIN in Cytoscape, the jActivesModules Cytoscape plugin was used to detect statistically significant differentially activated subnetworks (modules) at the 24 hpi and 48 hpi time points. The top five subnetworks at each time point were retained for downstream analyses and consisted of 198 genes in module 1 at 24 hpi (M1-24), 287 genes in M2-24, 272 genes in M3-24, 53 genes in M4-24, 171 genes in M5-24, 381 genes in M1-48, 330 genes in M2-48, 403 genes in M3-48, 371 genes in M4-48, and 399 genes in M5-48 (Supplementary Data 5). As an example, Fig. 4b shows the subnetwork of genes and gene interactions representing module 5 at 24 hpi (M5-24).

**Fig. 4.**
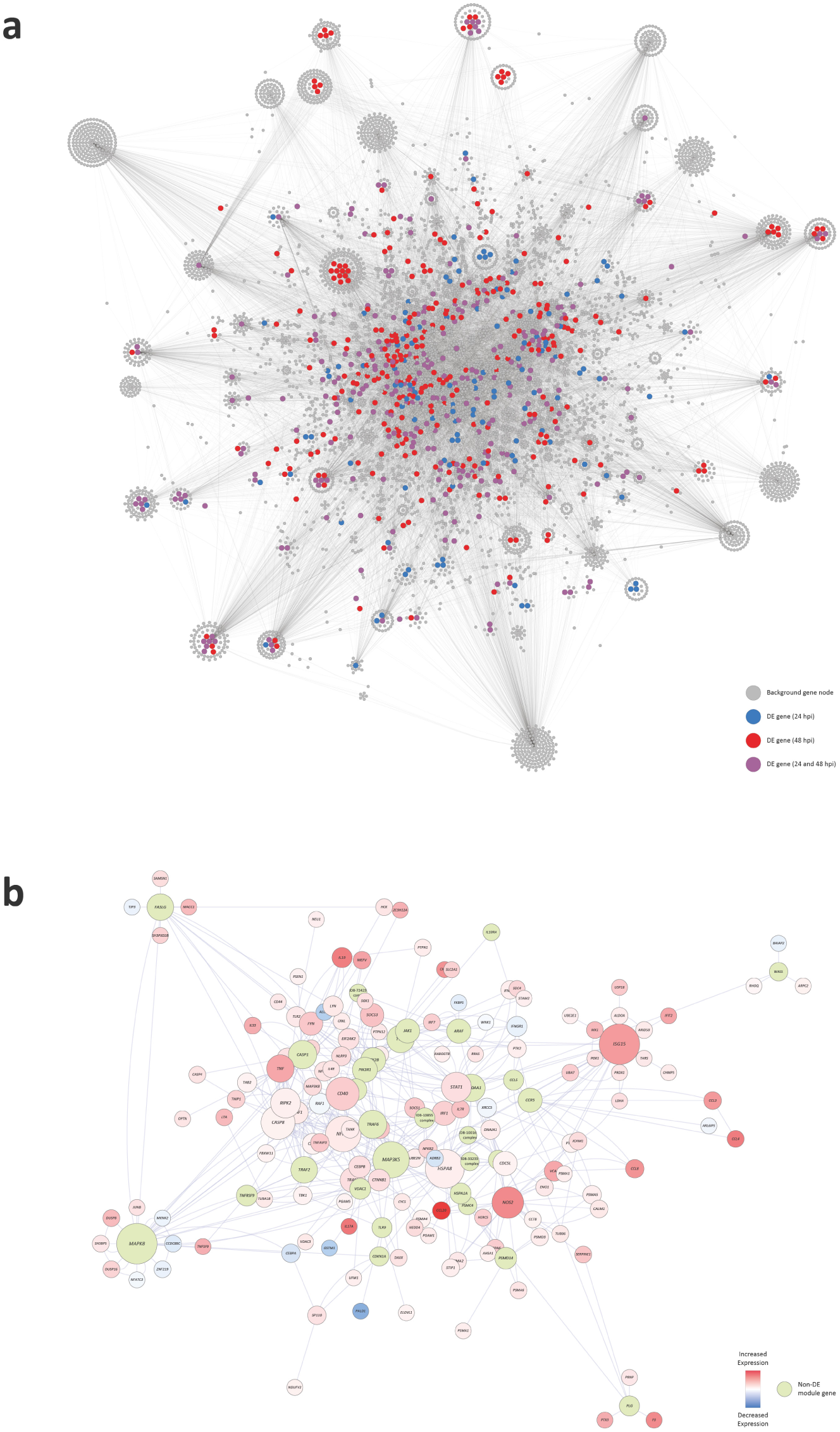
A large gene interaction network (GIN) with superimposed differentially expressed genes. **a** Cytoscape tuberculosis gene interaction network with superimposed differentially expressed genes at 24 and 48 hpi. **b** Example subnetwork showing module number 5 detected with the 24 hpi gene expression data.

The genes contained in the top five modules at 24 hpi and 48 hpi were filtered to remove duplicates and consolidated into two separate gene sets for GWAS integration with *gwinteR*. The consolidated gene sets for the top five modules at 24 hpi and 48 hpi contained 339 and 495 unique genes, respectively. There were 245 genes that overlapped between the two subnetwork gene sets for the two post-infection time points. The two differential expression network (DEN) *gwinteR* input gene sets (DEN-24 and DEN-48 – see Fig. 1) are also detailed in Supplementary Data 5.

### GWAS integration and identification of additional SNP–trait associations

The six gene sets generated from the three separate analyses of DE genes in bAM challenged with M. bovis at 24 hpi and 48 hpi are summarised in Table 2 and further detailed in Supplementary Data 2–4. In addition to these six putative functionally relevant gene sets, one hundred sets of 250 genes randomly sampled from the bovine genome were used for statistical context and comparison. These random gene sets (RAN) are detailed in Supplementary Data 6 (see also Fig. 1). The results from the integrative analyses using *gwinteR* with the DEG-24, DEG-48, CON-24, CON-48, DEN-24, DEN-48, and RAN gene sets are summarised graphically in Fig. 5 and detailed in Supplementary Data 7 and 8. Fig. 5a shows circular Manhattan plots with GWAS results (*P*_adj_. values) for each of the three breeds prior to data integration using *gwinteR*. Fig. 5b shows the *gwinteR* permuted *P*-values (*P*_perm_.) for each of the 10 genomic intervals used and for each of the six input gene sets plus the RAN gene set with *P*_adj_. < 0.10. Fig. 5c shows circular Manhattan plots with GWAS results post data integration using *gwinteR*. Inspection of Fig. 5b shows that, in terms of SNP enrichment (*P*_perm_. < 0.05), the integrative analyses using *gwinteR* were most effective for the HOFR breed group where the CON-24, CON-48, and DEG-48 input gene sets produced enriched SNPs across all 10 genomic ranges. In addition, the DEG-24 and DEN-24 input gene sets were effective for the HOFR breed across the ± 20 to 100 kb and ± 30 to 50 kb genomic ranges, respectively. In the case of the LIM breed, the DEG-48 input gene set produced enriched SNPs across all 10 genomic ranges, the CON-48 between ± 10 to 70 kb and the CON-24 at ± 10 kb. For the CHA breed, SNP enrichment using *gwinteR* was only observed for the CON-24 input gene set for the genomic interval between ± 10 to 40 kb.

**Table 2:**
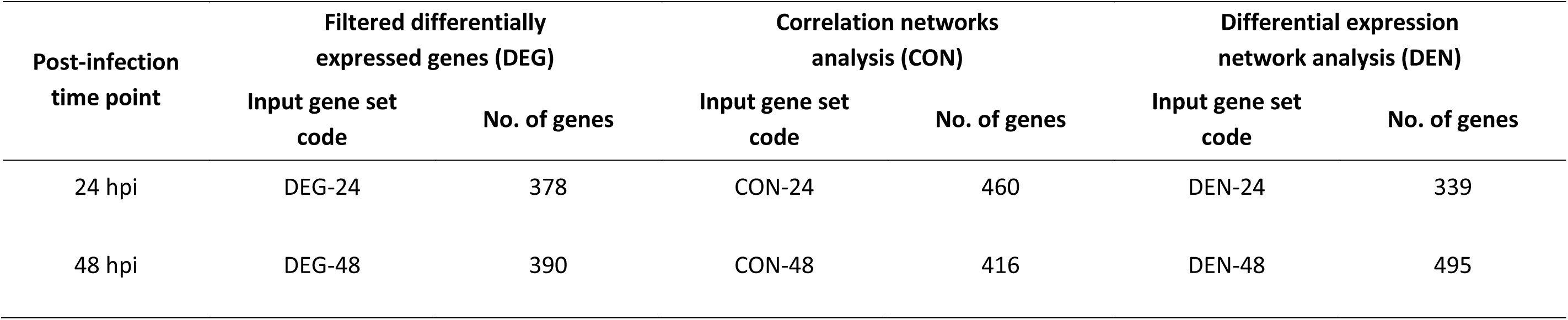
Six different input gene sets used for GWAS integration

**Fig. 5.**
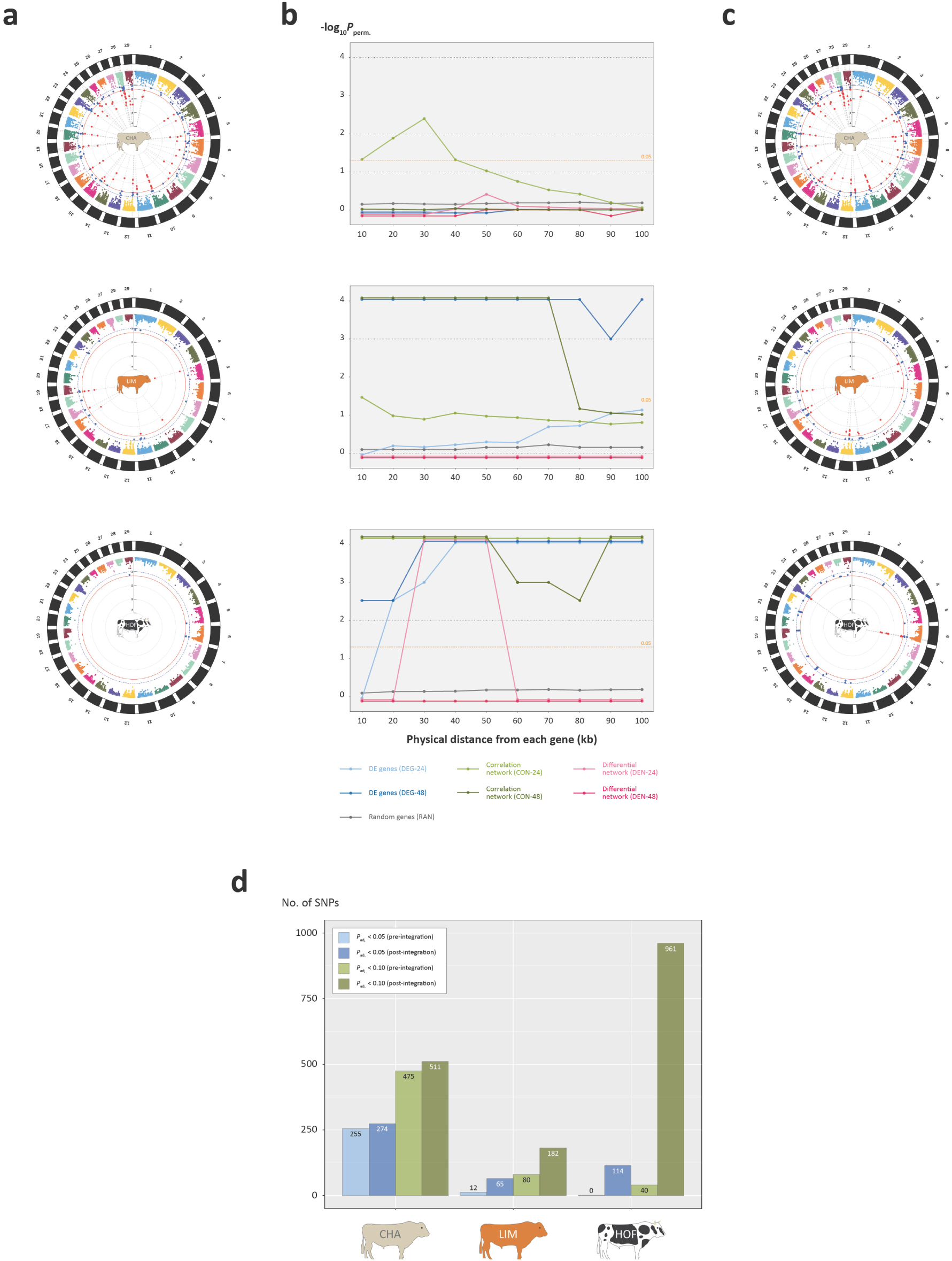
Integration of bAM functional genomics and GWAS data for resistance to *M. bovis* infection in three cattle breeds. **a** Circular Manhattan plots showing GWAS results pre-integration with blue and red data points indicating binned SNP clusters with FDR *P*_adj_. < 0.10 and < 0.05, respectively. **b** Line plots of permuted *P*-values across different genomic intervals for SNPs from six different input gene sets. **c** Histogram showing numbers of significant GWAS SNPs for the bTB resistance trait, pre-and post-enrichment for the three cattle breeds.

Fig. 5d summarises the numbers of statistically significant SNPs pre-and post-data integration, again with SNP enrichment being most evident for the HOFR breed with a 24-fold post-*gwinteR* SNP enrichment at *P*_adj_. < 0.10 from 40 to 961 SNPs. The SNP enrichments for the CHA and LIM breed were more modest, although there was a 2.3-fold enrichment at *P*_adj_. < 0.10 from 80 to 182 SNPs for the LIM breed. Inspection of Supplementary Data 8 reveals notable gene loci associated with enriched GWAS SNPs for the HOFR breed including the allograft inflammatory factor 1 gene (*AIF1*), which encodes a protein that promotes macrophage activation and proinflammatory activity (Liu *et al.* 2007); the ciliogenesis associated kinase 1 gene (*CILK1* aka *ICK*); *IL17A* and *IL7F*, which encode proinflammatory cytokines and contain polymorphisms in the human orthologs that have been associated with lower human TB disease incidence (Wang *et al.* 2016; Eskandari-Nasab *et al.* 2018); the integrin subunit beta 3 gene (*ITGB3*), which encodes a protein that has been shown to regulate matrix metalloproteinase secretion in pulmonary human TB (Brilha *et al.* 2017); the neuraminidase 1 gene (*NEU1*), which encodes a protein that regulates phagocytosis in macrophages (Seyrantepe *et al.* 2010); and the *TNF* gene that encodes TNF (aka TNF-α), a key cytokine for generation and maintenance of the granuloma, and where gene polymorphisms have been linked to resistance to *M. bovis* infection (Cheng *et al.* 2016). Gene loci associated with enriched GWAS SNPs for the CHA breed group also included the *CILK1* gene (aka *ICK*) and the Kelch repeat and BTB domain containing 3 gene (*KCNJ15*), which has been detected as an expression biomarker for human TB (Satproedprai *et al.* 2015); the T cell immune regulator 1, ATPase H+ transporting V0 subunit a3 gene (*TCIRG1*), a known antimycobacterial host defence gene that has been shown to be a key hub gene associated with IFN-γ stimulation of human macrophages (Steiger *et al.* 2016); and the Von Willebrand factor gene (*VWF*). Notable gene loci associated with enriched GWAS SNPs for the Limousin breed group included the cellular repressor of E1A stimulated genes 1 gene (*CREG1*), which encodes a regulator of core macrophage differentiation genes (Gautier *et al.* 2012); the desmoplakin gene (*DSP*), which increases in expression during *M. tuberculosis*-derived ESAT6-regulated transition of bone marrow-derived macrophages (BMDMs) into epithelioid macrophages (Lin *et al.* 2020); and the SP110 nuclear body protein gene (*SP110*), which encodes a protein that modulates growth of MTBC pathogens in macrophages and has been successfully exploited for genome editing of cattle to enhance resistance to *M. bovis* infection (Wu *et al.* 2015)

## Discussion

During the last decade, integrative genomics, multi-omics analyses and network biology have come to the fore as powerful strategies for exploring, dissecting and unpicking the complexities of the vertebrate immune system and immune responses to specific microbial pathogens (Schubert 2011; Kidd *et al.* 2014; Vodovotz *et al.* 2017; Hao *et al.* 2020). In the present study we have used these approaches to integrate transcriptomics data from a pivotal immune effector cell in *M. bovis* infection with high-resolution GWAS data for a bTB resistance trait. We developed a new computational tool, *gwinteR*, to enhance detection of QTLs by leveraging nominal SNP *P*-values from large GWAS data sets for resistance to infection by *M. bovis* in cattle. Three different integration strategies were employed with transcriptomics data from bAM infected with *M. bovis* across a 48-h time course. The first and most straightforward method was based on DE gene sets at 24 and 48 hpi with stringent fold-change and *P*-value thresholds as filtering criteria (DEG-24 and DEG-48). For the second method, a correlation network approach was used to identify subnetworks (modules) of co-expressed functional gene clusters from the bAM transcriptomics data at the two post-infection time points (CON-24 and CON-48). The third method was also network-based but took advantage of the extensive scientific literature and curated biomolecular data for mycobacterial infections and tuberculosis disease. For this approach, a base GIN was constructed and functional modules containing overlaid differentially expressed genes were identified to provide two post-infection input gene sets (DEN-24 and DEN-48) for downstream data integration.

Integration of these six input gene sets with bTB GWAS data from three different cattle breeds (CHA, LIM and HOFR) revealed substantial differences among the three methods in their capacity to detect additional QTLs for a *M. bovis* infection resistance trait in these particular GWAS data sets. For example, the correlation network approach was the only method that enriched SNPs for the CHA breed and that worked for at least one bAM post-infection time point across all three breeds (see Fig. 5). Surprisingly, perhaps, the functional modules obtained using the GIN differential network (DEN) approach—the most complex method to implement—produced the least effective input gene sets for prioritising additional SNPs from the GWAS data set. This method proved effective only for the DEN-24 input gene set with the HOF breed and then only for the ±30, ±40, and ±50 kb genomic intervals. Conversely, the simplest method based on DE genes at 24 and 48 hpi enriched SNPs for both the LIM and HOF breed groups, with the DEG-48 input gene set being the most effective (Fig. 5).

In summary, across the two different bAM infection time points and the three breeds, the CON method enriched 970 SNPs, the DEG method enriched 163 SNPs and the DEN method enriched only 11 SNPs (Supplementary Data 8). Although other factors such as linkage disequilibrium need to be considered in interpreting these differences, it is reasonable to hypothesise that GWAS integration using the correlation network approach is more sensitive to regulatory genomic variants that alter expression of co-ordinately regulated protein components of the alveolar macrophage pathways and processes underpinning host-pathogen interaction for the early stages of intracellular MTBC infection (Weiss & Schaible 2015; Kaufmann & Dorhoi 2016; Schorey & Schlesinger 2016). Fig. 6 illustrates this using the example of SNPs within and proximal to gene loci associated with PI3K/AKT signalling, which based on IPA data mining for bAM DE genes was a highly activated pathway, particularly at 24 hpi. In this regard, previous work has shown that macrophage PI3K/AKT signalling is key to a range of cellular processes associated with host-pathogen interaction in MTBC infections, including modulation of cell death pathways, manipulation of signalling downstream of TLRs, and initiation of granuloma formation (Maiti *et al.* 2001; A *et al.* 2012; Cho *et al.* 2013; Liu *et al.* 2016; Brace *et al.* 2017; Yang *et al.* 2018). We have also recently demonstrated that genes encoding protein products embedded in the PI3K/AKT pathway are primary targets for chromatin modifications that substantially alter bAM gene expression in response to *M. bovis* infection (Hall *et al.* 2020).

**Fig. 6.**
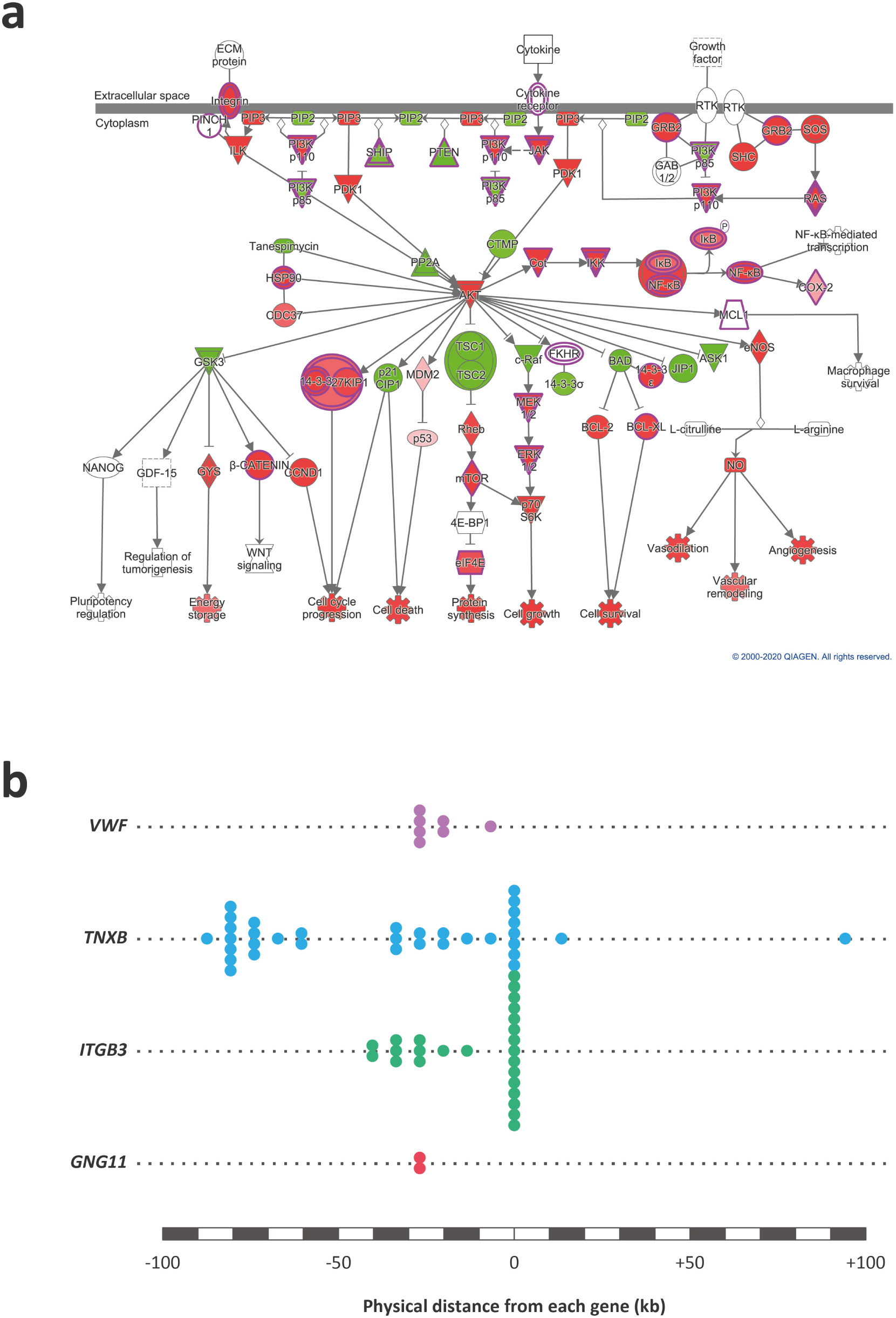
The PI3K/AKT signalling pathway and genomic variants within associated genes. **a** The IPA-predicted activation state of the PI3K/AKT signalling pathway with red and blue colours indicating increased and decreased activity of pathway components, respectively. **b** Schematic of enriched bTB disease resistance GWAS SNPs in four genes associated with the PI3K/AKT signalling pathway.

There were also notable breed differences in the effectiveness of the three methods used for multi-omics data integration and enrichment of GWAS SNPs. The original GWAS data set for the HOF breed was the least powered of the three breeds in terms of both sample size (*n* = 1,502 sires) and genetic markers (12,740,315 genome-wide SNPs). This is reflected in the relatively small number of GWAS SNPs that were detected pre-integration for the HOF breed: 40 SNPs at *P*_adj_. < 0.10 compared to 475 for the CHA breed and 80 for the LIM breed (see Fig. 5). However, the SNP enrichment post-integration was markedly more effective for the HOF breed with a 24-fold increase from 40 to 961 SNPs (*P*_adj_. < 0.10) compared to 1.08-fold (475 to 511 SNPs; *P*_adj_. < 0.10) and 2.28-fold (80 to 182 SNPs; *P*_adj_. < 0.10) for the CHA and LIM breeds, respectively (Fig. 5). In addition, a total of 48 genes were captured by the enriched GWAS SNPs for the HOF breed compared to 20 and 16 genes for the CHA and LIM breeds, respectively (Supplementary Data 8).

Regarding the superior performance of the HOF breed GWAS for multi-omics integration, it is perhaps noteworthy that the *M. bovis* infection transcriptomics data set was generated using bAM from this population (Nalpas *et al.* 2015). Most of the SNPs (*n* = 829, 88%) and genes (*n* = 36, 67%) obtained post-integration in the HOF breed were detected using the correlation co-expression network method, which, again, is likely caused by enrichment of genomic regulatory variants that modulate expression of genes associated with alveolar macrophage pathways and processes critical to early host-pathogen interaction. This pattern of variation may also reflect the polygenic architecture of *M. bovis* infection resistance in domestic cattle (Raphaka *et al.* 2017; Tsairidou *et al.* 2018a; Tsairidou *et al.* 2018b; Ring *et al.* 2019).

Integrative multi-omics approaches to data integration are now widely used to explore and dissect the genomic architecture and physiological basis of complex traits in domestic livestock, including network-based methods to integrate functional genomics and GWAS data (Canovas *et al.* 2014; Fang *et al.* 2017; Cai *et al.* 2018; Fang *et al.* 2018; Deng *et al.* 2019; Yan *et al.* 2020). However, to the best of our knowledge, this study is the first that uses network biology to systematically combine transcriptomics data from *M. bovis*-infected macrophages with GWAS data for *M. bovis* infection resistance in cattle. Therefore, it provides a novel framework for integrative genomics studies of complex infectious disease resistance traits in livestock, particularly those involving other intracellular bacterial pathogens such as *Brucella abortus*, *Mycobacterium avium* subsp. *paratuberculosis* and *Salmonella enterica*. The work is also relevant to development of methods for integrative analyses of outputs from the Functional Annotation of Animal Genomics (FAANG) initiative (Giuffra *et al.* 2019) and for identification and prioritization of targets for genome editing to enhance resistance to infection in domestic livestock species (Bishop & Van Eenennaam 2020). The results from this study may also inform genome-enabled breeding programmes for resistance to *M. bovis* infection in production cattle populations (Banos *et al.* 2017; Tsairidou *et al.* 2018a). Finally, this integrative multi-omics approach could also be used to combine relevant functional genomics and GWAS data sets to improve knowledge of innate immune responses and establishment of infection in human TB caused by *M. tuberculosis*.

## Data availability

The RNA-seq data set was generated by the authors and can be obtained from the NCBI Gene Expression Omnibus (GEO): accession number GSE62506. GWAS summary statistics data were obtained from a published study that provides additional information about sequence and genotype data availability (Ring *et al.* 2019).

## Code availability

Complete pipeline: https://github.com/ThomasHall1688/Bovine_multi-omic_integration

gwinteR: https://github.com/ThomasHall1688/gwinteR

DESeq2: https://github.com/mikelove/DESeq2

DGCA: https://github.com/andymckenzie/DGCA

MEGENA: https://github.com/cran/MEGENA

Cattle genome assembly and annotation: www.ncbi.nlm.nih.gov/datasets/genomes/?txid=9913

## Acknowledgements

This study was supported by Science Foundation Ireland (SFI) Investigator Programme Awards to D.E.M. and S.V.G. (grant nos. SFI/08/IN.1/B2038 and SFI/15/IA/3154); a Department of Agriculture, Food and the Marine (DAFM) project award to D.E.M (TARGET-TB; grant no. 17/RD/US-ROI/52); and a European Union Framework 7 project grant to D.E.M. (no: KBBE-211602-MACROSYS). The funding agencies had no role in the study design, collection, analysis, and interpretation of data, and no role in writing the manuscript.

## Author contributions

T.J.H, M.P.M., K.E.K., S.V.G and D.E.M. conceived and designed the study. J.A.B. performed experimental work. S.C.R. and D.P.B. provided cattle genotype and other genomic data. T.J.H., M.P.M., G.J.M., K.E.K. and C.N.C. performed bioinformatics and computational analyses. T.J.H., C.N.C. and D.E.M. wrote and prepared the manuscript and figures. All authors read and approved the final manuscript.

## Supplementary Information

**Supplementary Figure 1.** Principal component analysis (PCA) plots for individual animal bAM gene expression data at **a** 2 hpi, **b** 6 hpi, **c** 24 hpi, and **d** 48 hpi.

**Supplementary Figure 2.** Gene ontology (GO) enrichment for functional modules identified from the differential co-expression correlation network generated from *M. bovis*-infected bAM gene expression at 24 hpi.

**Supplementary Figure 3.** Gene ontology (GO) enrichment for functional modules identified from the differential co-expression correlation network generated from *M. bovis*-infected bAM gene expression at 48 hpi.

**Supplementary Data 1.** RNA-seq statistics and results.

**Supplementary Data 2.** Differentially expressed genes across the infection time course and DEG-24 and DEG-48 input gene sets for GWAS integration.

**Supplementary Data 3.** Enriched IPA canonical pathways at 24 and 48 hpi.

**Supplementary Data 4.** Outputs from correlation network analyses using DGCA with MEGENA and CON-24 and CON-48 input gene sets for GWAS integration.

**Supplementary Data 5.** Outputs from differential network analyses using GeneCards^®^, InnateDB with Cytoscape/jActiveModules and DEN-24 and DEN-48 input gene sets for GWAS integration.

**Supplementary Data 6.** 100 random (RAN) input gene sets used for GWAS integration.

**Supplementary Data 7.** Functional genomics and GWAS integration results for Charolais (CHA), Limousin (LIM) and Holstein-Friesian (HOF).

**Supplementary Data 8.** GWAS prioritised SNP results for Charolais (CHA), Limousin (LIM) and Holstein-Friesian (HOF).

