## Supplementary Figures for "Integrative genomics of the mammalian alveolar macrophage response to intracellular mycobacteria"

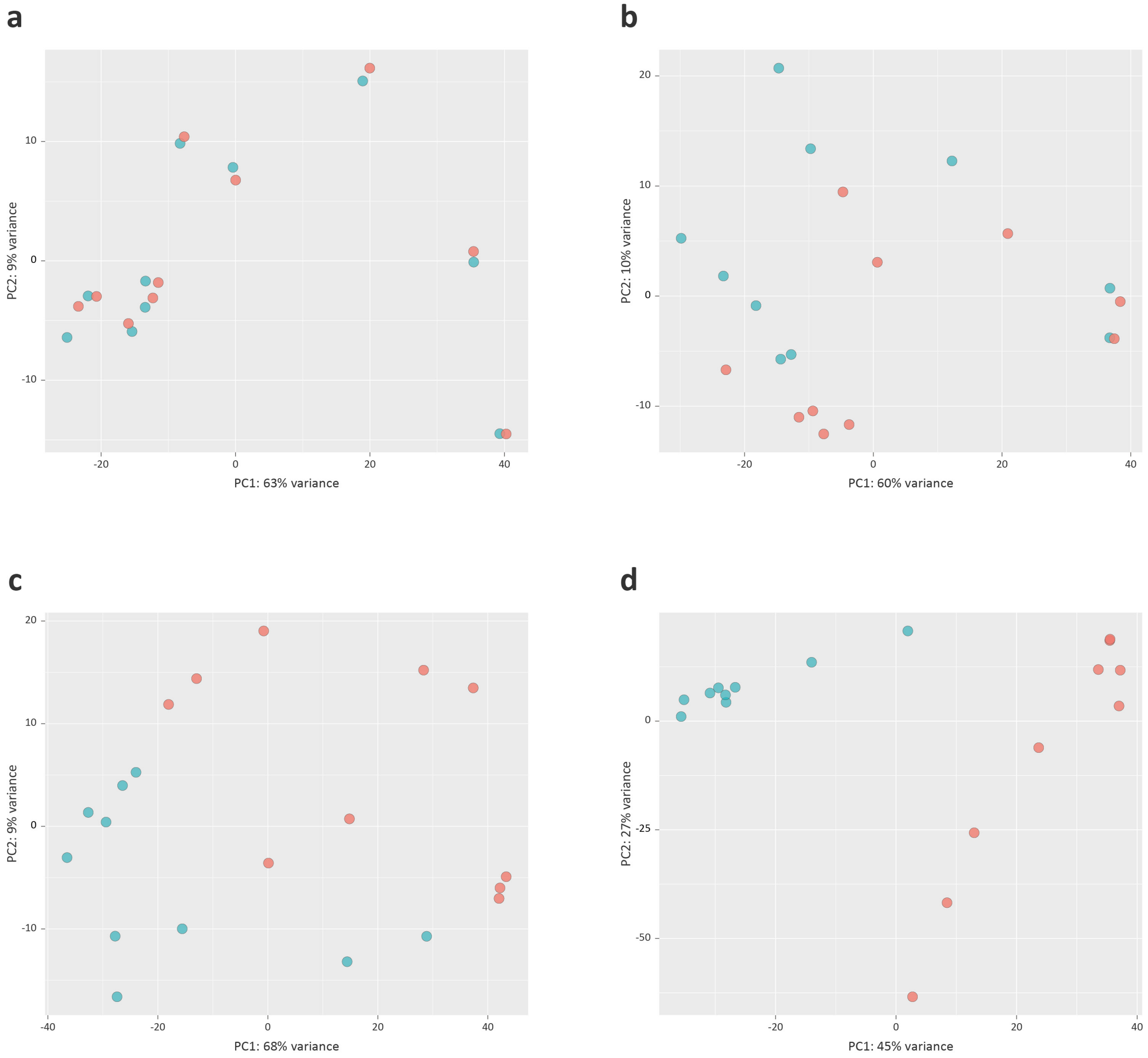


**Supplementary Figure 1.** Principal component analysis (PCA) plots for individual animal bAM gene expression data at **a** 2 hpi, **b** 6 hpi, **c** 24 hpi, and **d** 48 hpi. Red indicates *M. bovis*-infected animals and blue indicates non-infected control animals.

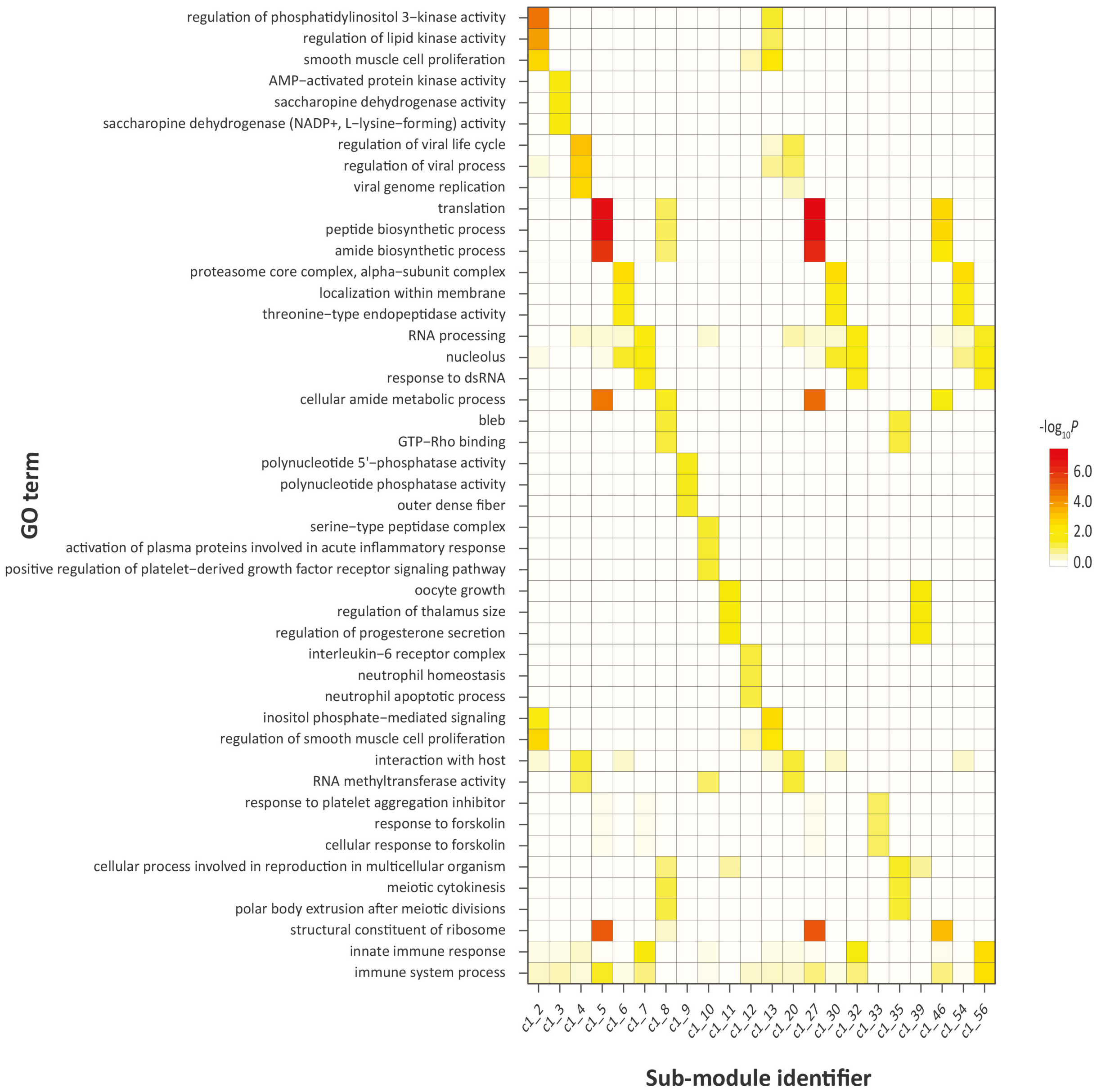


**Supplementary Figure 2.** Gene ontology (GO) enrichment for functional modules identified from the differential co-expression correlation network generated from *M. bovis*-infected bAM gene expression at 24 hpi.

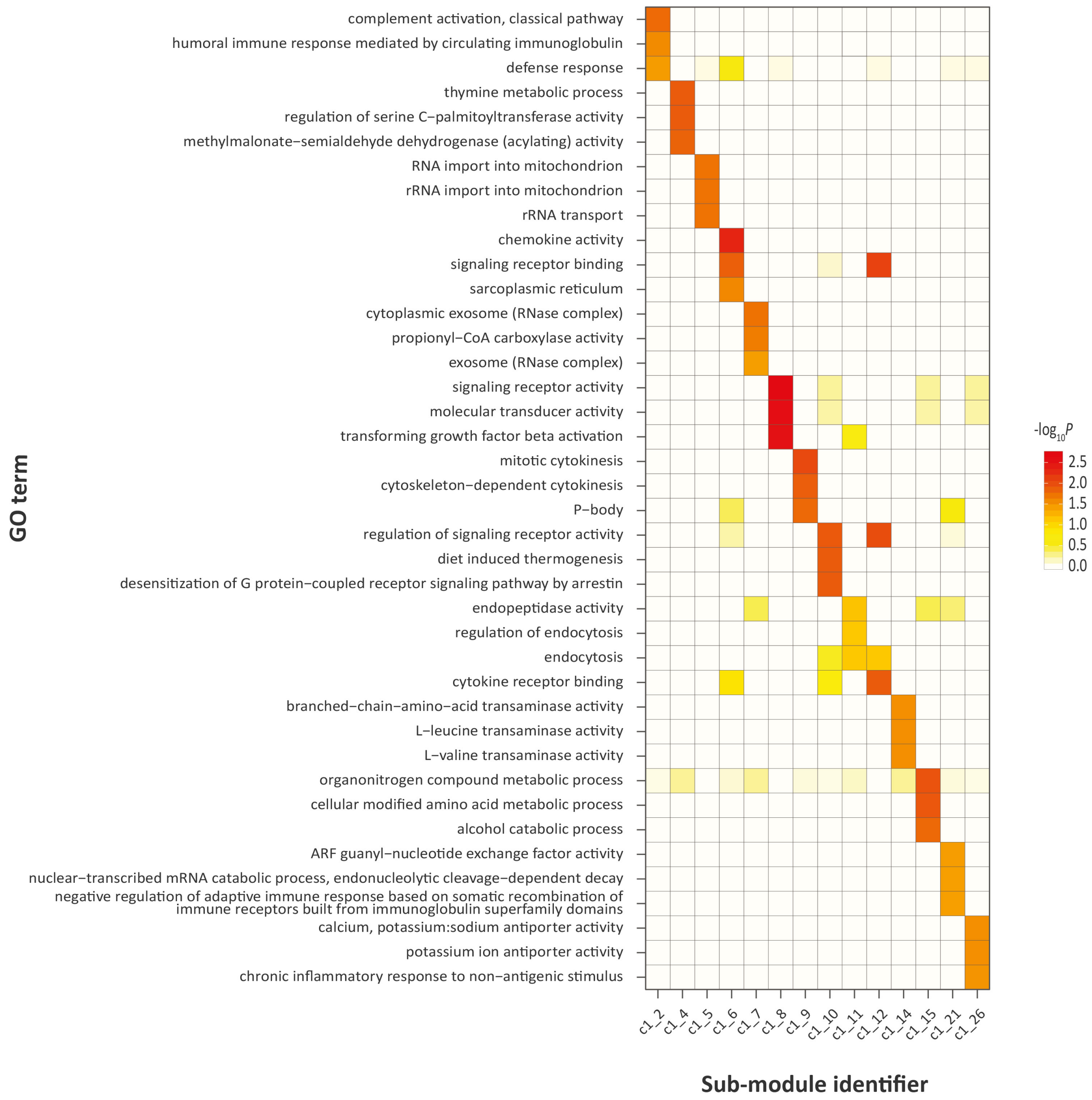


**Supplementary Figure 3.** Gene ontology (GO) enrichment for functional modules identified from the differential co-expression correlation network generated from *M. bovis*-infected bAM gene expression at 48 hpi.
